# Explicit description of viral capsid subunit shapes by unfolding dihedrons

**DOI:** 10.1101/2024.07.29.605617

**Authors:** Ryuya Toyooka, Seri Nishimoto, Tomoya Tendo, Takashi Horiyama, Tomohiro Tachi, Yasuhiro Matsunaga

**Affiliations:** Department of General Systems Studies, The University of Tokyo, Tokyo; Faculty of Information Science and Technology, Hokkaido University, Sapporo; Graduate School of Science and Engineering, Saitama University, Saitama

**Keywords:** Quasi-equivalence, Caspar-Klug theory, Spherical tiling, Dihedron, Molecular docking

## Abstract

Viral capsid assembly and the design of capsid-based nanocontainers critically depend on understanding the shapes and interfaces of constituent protein subunits. However, a comprehensive framework for characterizing these features is still lacking. Here, we introduce a novel approach based on spherical tiling theory that explicitly describes the 2D shapes and interfaces of subunits in icosahedral capsids. Our method unfolds spherical dihedrons defined by icosahedral symmetry axes, enabling systematic characterization of all possible subunit geometries. Applying this framework to real *T* = 1 capsid structures reveals distinct interface groups within this single classification, with variations in interaction patterns around 3-fold and 5-fold symmetry axes. We validate our classification through molecular docking simulations, demonstrating its consistency with physical subunit interactions. This analysis suggests different assembly pathways for capsid nucleation. Our general framework is applicable to other triangular numbers, paving the way for broader studies in structural virology and nanomaterial design.

## Introduction

Viral capsids are assemblies of proteins that encapsulate and protect the viral genome. Many spherical viral capsids adopt icosahedral structures, which is fully characterized by 60 symmetry operations. To elucidate the mechanism of self-assembly of molecules^1,2^ and also the rational design of capsid-based nanocontainers^3–6^, it is important to understand how icosahedral symmetry imposes geometric constraints on the interaction patterns between subunit proteins.

The Caspar-Klug (CK) theory is currently used as a major tool for classifying capsid structures^7^. The theory explains how capsids can be formed from different numbers of subunits, resulting in various sizes^8^. In the CK theory, protein subunits are modeled using a hexagonal network of subunits on a plane according to the *p*6 wallpaper group, whose 6-fold symmetric interface on the plane is regarded as quasi-equivalent to that of the 5-fold symmetry axis on the icosahedron. The CK theory systematically describes the size and number of subunits by subdividing the triangular region into multiple triangles using two integers (*h, k*) and the triangulation (*T)* numbers. On the other hand, the CK theory cannot directly address questions on what the necessary shapes of subunits and interfaces for self-assembling icosahedral capsids are. This question is not only essential for the rational design of capsids of subunit proteins but also for understanding the possible formation pathways during the self-assembling process. To answer this question, an explicit description or characterization of all possible subunit shapes or interfaces would be needed.

Thus far, several contributions have directly or indirectly focused on characterizing the shapes of subunits in viral capsids from viewpoints beyond the CK theory^9–15^. These studies include affine extensions to describe the icosahedral groups and virus capsids^10,12^, and a framework to investigate the locations of protrusions^13^. Twarock introduced non-triangle tiles instead of the CK theory’s triangular tiles to characterize Simian Virus 40 and L-A virus capsids^9^. Raguram et al. also developed a general polyhedral framework to describe virus capsid structures, employing pentagonal subunits, accounting for intrinsic capsid chirality^14^. Twarock and Luque recently elegantly extended the CK theory by using non-CK tiles (Archimedean tiles and their duals) to describe capsid structures that fall outside the CK description^15^. While these works significantly broadened the spectrum of describable capsid geometries beyond the original CK theory, the subunit shapes that can be systematically handled are still confined to specific geometry forms, such as typical pentagonal, hexagonal, and triangular tiles, as well as rhombs, kites, and florets in the dual representations.

The contribution of this study is to develop a framework for describing the shapes and interfaces of subunit proteins based a novel representation using the tiling theory on spherical surfaces. The idea is based on the unfolding of dihedrons with the triangular shape used in the CK theory. Our representation can account for all possible 2D shapes and interfaces of subunit proteins of icosahedral capsids. Using the proposed representation, we classify the icosahedral structures according to the shapes and interfaces of their subunits in the *T* = 1 group. Then, we show that there are distinct interface types within the same *T* = 1 group, with variations in interface areas around the 5-fold, 3-fold, and 2-fold symmetry axes. We propose these interface areas as a governing factor to characterize the interactions between subunits. Finally, we perform pairwise docking simulations of two subunit proteins to validate that the favored interface computed from the docking score and the predicted interface from the classification are consistent with each other. These results imply that there are multiple types of assembling interactions in capsid structures and a strategy for creating a nucleus in the self-assembly process.

## Results

### Dihedron-unfolding model based on spherical tiling

We here focus on *T* = 1 tiling for simplicity, while our approach is general and can straightforwardly extend to *T* ≠ 1 cases (see Supplementary Fig. 1 and text for extension). In *T* = 1 tiling, 60 subunits are assembled according to chiral icosahedral symmetry. Subunit shapes can be generated by creating the fundamental figure that makes up 1/60 of the sphere (Fig. 1 Middle). The boundary of such a figure can be segmented into at most three pairs of identical curves copied by rotations of 180^°^, 120^°^, and 72^°^ about the 2-, 3- and 5-fold symmetry axes, respectively. Consider that the tile is made of a thin sheet; then stitching the identical boundary curves together yields a dihedral shape, precisely, a double-covering of the spherical triangles formed between the three axes of symmetries (1/120 of the sphere) (Fig. 1 Left). Any tile figure can be obtained by its inverse process; i.e., we start from the dihedron, cut it along an arbitrary curve, and unfold it into a spherical tile (Supplementary Movie 1). To unfold the polyhedron with three vertices into a topological disk, the dihedron must be cut along a tree graph with three leaves at the vertices; specifically, the graph is a Y-shape whose three ends correspond to the vertices. To focus on the contact relationship, the cut graph is simplified to three geodesics connected at the *junction point*, i.e., the degree-3 vertex and three vertices (Fig. 1 Right). Then, the unfolding is identified by the choice of orientation of the dihedron and the location of the junction point of the cut tree. The former orientation corresponds to the choice of either a left- or right-handed triangle as the basis for the dihedron (Here, the triangle with vertices 2, 3, and 5 in counterclockwise order is called the right-handed). We construct a polygonal tile by alternately connecting the three axes and the three mirror reflections of the junction point with respect to the edges of the spherical triangle.

**Figure 1.**
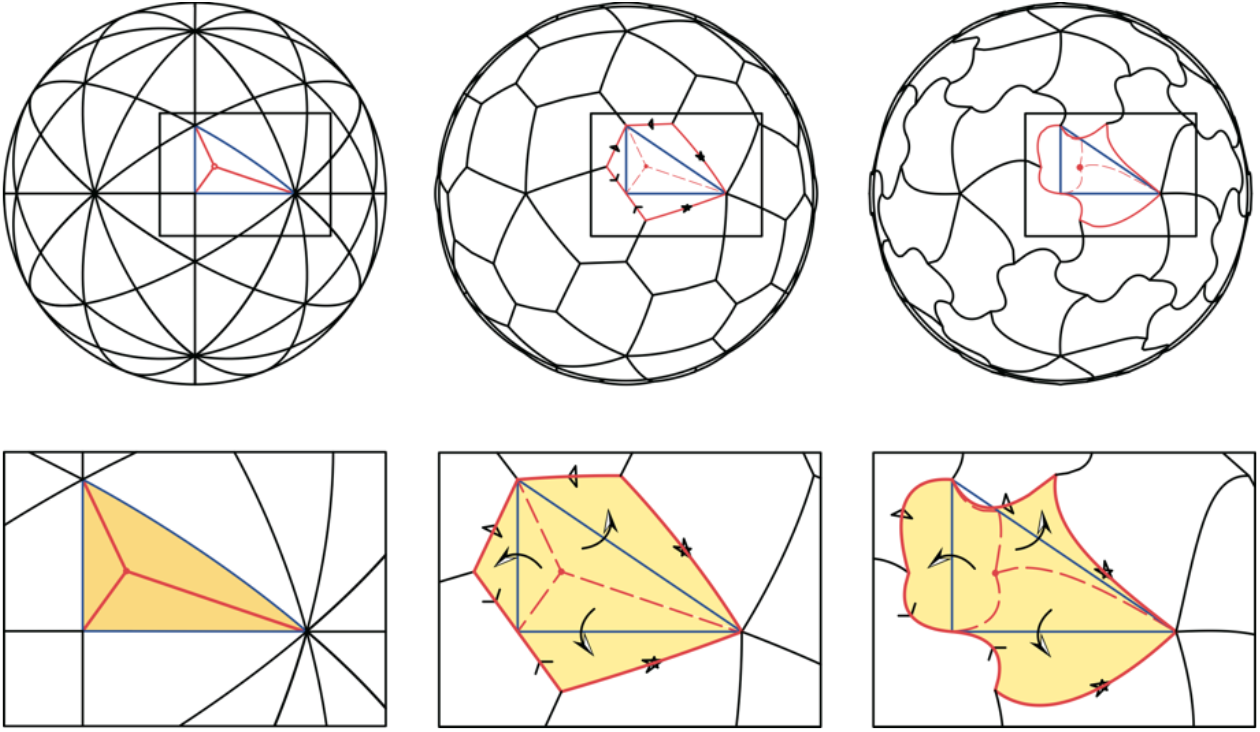
Dihedron unfolding and the tile shapes. Left: dihedron, i.e., double covering of 1/120 of the sphere and the cut tree. Middle: the unfolding of the dihedron forming a tile using the cut tree. Right: a cut tree simplified using three geodesics.

This dihedron-unfolding model provides two-parameter representation of the contact between subunits arranged in *T* = 1 group. Figure 2 shows the different tile shapes and connectivities corresponding to different positions of the junction point and the orientation. Note that the junction point can be placed outside the triangle. Even in this case, the shape of the tile can be defined similarly by mirror reflection with respect to each edge, resulting in a concave tile (Fig. 2 Top Left and Top Right. Supplementary Movies 2 and 3). The position of the junction point is restricted to the highlighted area illustrated in Fig. 2; if the junction point is outside this region, the unfolded tile self-intersects (i.e., mirror-reflected edges intersect with each other).

**Figure 2.**
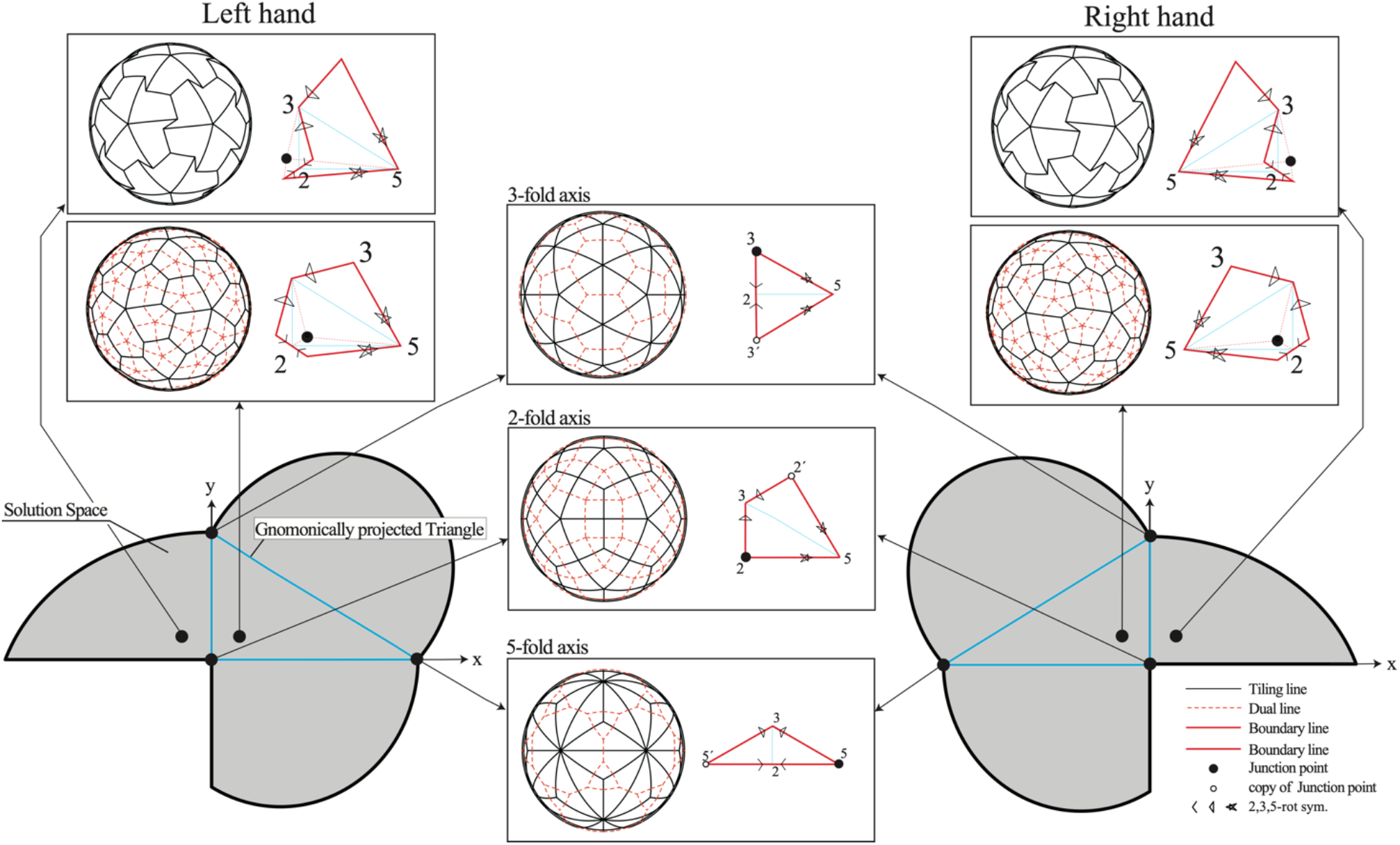
Dihedron-unfolding model for tile shapes and contacts. For each digon, a gray range shows the permitted positions of the junction point. If the junction point is outside the gray range, the unfolded figure self-intersects. In the generic case, the resulting tile has chirality. The left and right triangles correspond to the left- and right-handed orientation of the dihedron. If the junction point is outside the triangle, the tile is concave. In a degenerate case, the tiling is achiral when the junction point lies on the symmetry axis.

The connectivity between the tiles can be represented by the polyhedral graph and its dual graph. In the dual graph, the node corresponds to the subunit, and the edge connecting the two nodes represents the connectivity between the subunits. For visualization purposes, we use gnomonic projection, which represents geodesic lines as straight lines, to specify the junction point as a point on a projected plane; here, we set that the projection plane is perpendicular to the 2-fold symmetric axis (*z*-axis in Fig. 2).

In a *generic case*, i.e., when the junction point is not on the symmetry axis, each tile is pentagonal. Thus, each subunit is surrounded by five neighboring subunits: one copy by 2-fold symmetry, two copies by 3-fold symmetry, and two copies by 5-fold symmetry. The polyhedral graph of the tiling is homeomorphic to the pentagonal hexecontahedron; thus, its dual is homeomorphic to the snub dodecahedron. This case includes the pentagonal tiles investigated by Raguram et al^14^ in their study to develop a general polyhedral framework. As already discussed by Raguram et al.^14^, the pentagonal hexecontahedron and snub dodecahedron have chirality, which depends on the choice of orientation when unfolding the dihedron. The left- and right-handed tilings (Laevo and Dextro, respectively) result in left- and right-handed pentagonal hexacontahedral graphs, respectively.

In *degenerate cases*, we locate the junction point on the 2-, 3-, and 5-fold symmetry axes. When the junction point is on the 2-fold symmetry axis, we obtain quadrangular tilings homeomorphic to the deltoidal hexecontahedron; thus, its dual is homeomorphic to the rhombicosidodecahedron. Each subunit is surrounded by four neighboring subunits: 2 copies by 3-fold symmetry and 2 copies by 5-fold symmetry. This case includes the kite-like tiles, as a special case, studied by Twarock and Luque^15^ for the analysis of Tobacco ringspot virus^16^ using non-CK tiles (Archimedean tiles and their duals). When the junction point is on the 3-fold symmetry axis, we obtain triangular tilings homeomorphic to the pentakis dodecahedron; thus, its dual is homeomorphic to the truncated icosahedron. Each subunit is surrounded by three neighboring subunits: 1 copy by 2-fold symmetry and 2 copies by 5-fold symmetry. When the junction point is on the 5-fold symmetry axis, we obtain triangular tilings homeomorphic to the triakis icosahedron; thus, its dual is homeomorphic to the truncated dodecahedron. Each subunit is surrounded by three neighboring subunits: 1 copy by 2-fold symmetry and 2 copies by 3-fold symmetry.

### Characterization of real capsid structures

We fitted our dihedron-unfolding model to experimental capsid structures of *T* = 1 to classify real structures. Here, by fitting the model to real structures, we inversely estimated the left- or right-handed orientation and location of the junction point for each capsid. We first determined left- or right-handed orientations based on the observation that the boundary of the tile passes through all three axes. So, we chose the orientation such that the maximum of the shortest distances from the axes to the atom positions (see Fig. 3a for the interpretation of the criteria) are smaller. Then, we calculated the fitness (measured by the Dice coefficient) between the real structures (represented by the circles with the effective radii of amino acids centered at their Cα atom positions of subunit protein) and the tile in the gnomonic projection and maximized the fitness using a genetic algorithm (see Methods for details).

**Figure 3.**
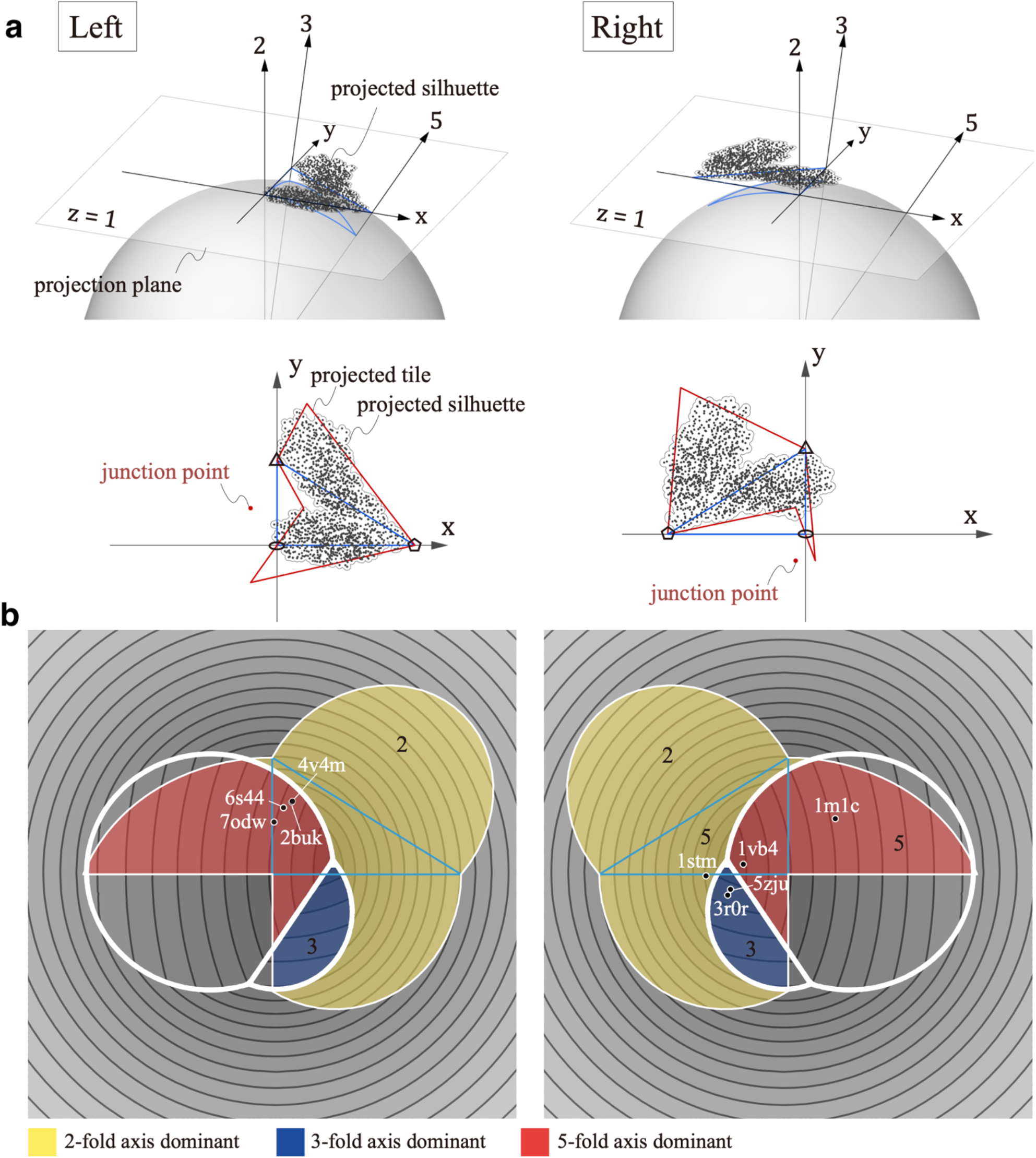
Fitting of real capsid structures by the dihedral unfolding framework. **a**. Gnomic projection of the Cα atom positions on an icosahedral capsid. The location of junction point that maximizes the Dice coefficient between the model and real shapes is searched on the projected plane. **b**. The orientations and the locations of the junction points fitted to real capsid structures. Highlighted region shows where the junction point does not induce self-intersection. The region is colored according to three domains governed by contact with respect to the 2- (yellow), 3- (blue), and 5-fold (red) axes.

Figure 3 illustrates the computed orientations and locations of the junction points for the real capsid structures. We used the capsid structures of *T* = 1 number taken from Protein Data Bank (PDB) whose PDB IDs are 2BUK^17^, 4V4M^18^, 6S44^19^, 7ODW^20^, 3R0R^21^, 5ZJU^22^, 1STM^23^, and 1VB4^24^. 2BUK and 4V4M belong to the satellite tobacco necrosis virus, both sharing identical sequences (structures are slightly different due to experimental conditions). Also, both 3R0R and 5ZJU belong to the porcine circovirus 2, albeit with slightly different sequences. This selection of redundant pairs aimed to validate the robustness of our fitting procedure. The other structures have different origins and sequences with each other: the faba bean necrotic stunt virus (6S44), a model of the Haliangium ochraceum encapsulin (7ODW), and the sesbania mosaic virus deletion mutant (1VB4). the structure of 1M1C^25^, classified as *T* = 2 comprising a 120-homomer, was also included. In this capsid, a neighboring subunit pair (dimer) was treated as a single tile in the fitting process. For details of the structural data, see Methods and Supplementary Fig. 2.

In Fig. 3b, all capsids fall into generic types, where 2BUK, 6S44, 7ODW, 4V4M are left-handed and 1VB4, 1STM, 5ZJU, 3R0R, and 1M1C are right-handed. In the figure, we provide a simple metric based on the lengths of shared edges between neighboring capsids copied by the 2-, 3- and 5-fold axes to evaluate the contacts between adjacents numerically. These lengths are the distances from the axis for 3- and 5-fold axes, while twice the distance is used for the 2-fold axis, as we consider the edges active in a face-to-face manner between a pair of subunits. As the longer contact edge is associated with the strong interaction between the subunits, the categorization with this measure suggests the strength of pairwise interaction between the subunits, which we verify in the following section. The contour plot shows the maximum value of the distances; this decomposes the region into three domains governed by contact with respect to the 2-, 3-, and 5-fold axes.

The figure shows that the 2BUK and 4V4M subunits, which have sequences identical to each other, possess the longest contact edge between their copies along the 5-fold axis in the left-handed. In contrast, the other twin subunits, 3R0R and 5ZJU (with almost identical sequences to each other), exhibit the longest contact edge around the 3-fold axis, characterized by a different chirality (the right-handed). Interestingly, other subunits also cluster around the same regions as these twin subunits, namely the left-handed 5-fold axis and the right-handed 3-fold axis. Since there is no physical reason to prefer any specific chirality for subunit interactions, the observed tendency for these junction points to cluster would be evolutionarily coincidental. Another intriguing observation is that almost all junction points do not distribute around regions governed by the 2-fold axis (except for 1STM). The 2-fold symmetry that interacts in a face-to-face manner is evolutionarily easier to optimize compared to other symmetry axes^26^. In fact, it is well known that structures stabilized around the twofold axis are common in ordinary dimers^27^. On the other hand, in the case of capsids, symmetries other than the 2-fold may be more important for creating a kinetic nucleus for growth toward full capsid formation.

### Pairwise docking simulation of subunit proteins

In order to investigate whether our classification of subunit contacts is consistent with physical interactions, we conducted rigid-body docking simulations of paired subunits for the experimental structures. In the rigid-body docking simulations, the structure of the subunit protein is treated as a rigid body, and no conformational changes are considered. We here used ZDOCK^28,29^ for the simulation. In ZDOCK, physical and statistically derived interaction energies are approximately calculated with regular grids and docking poses (translations and orientations) with high docking scores are exhaustively searched.

Figure 4 shows the results of the docking simulations. Here, to relate the docking poses to the symmetry axes of capsid, we calculated the screw axis^30^ for individual docking structure of homo-dimer. The screw axis is an axis that describes a rigid body motion (translation and rotation) to superimpose one monomer to the other one (Supplementary Fig. 5). The screw axis and translation and rotation around that axis were determined by using the Rodriguez equations (see Methods). The maximum scores of docking scores and the minimum values of root mean square deviation (RMSD) from the experimental structure are shown as heatmaps in the space of rotations and translations of the screw axes.

**Figure 4.**
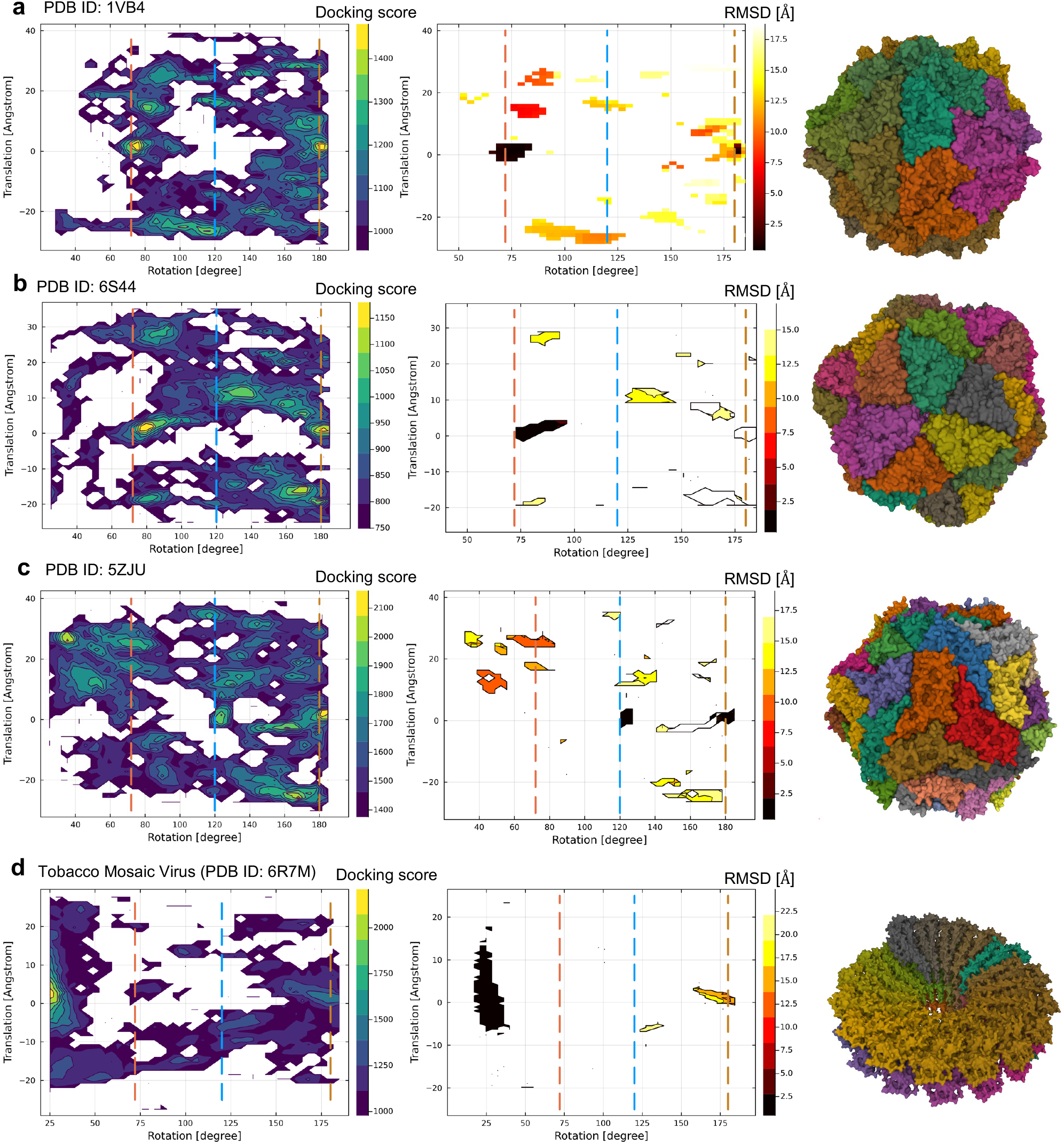
Results of rigid body docking simulations for a pair of subunits, **a**. PDB ID: 1VB4, **b**. 6S44, **c**. 5ZJU, and **d**. 6R7M (Tabacco mosaic virus). Left: heatmap of the maximum docking scores of docked poses in the space of rotation and translation of detected screw motions. Middle: heatmap of the minimum root mean square displacements of docked poses from the experimental capsid structure. Dashed lines indicate the rotation positions of 72^°^ (red), 120^°^ (blue), and 180^°^ (ocher), which correspond to the 5-fold, 3-fold, and 2-fold axes, respectively. Right: the experimental capsid and non-capsid structures. Different subunits are represented by different colors.

The figure shows that 1VB4 (classifiled as right-handed and 5-fold in our dihedron-unfolding model) and 6S44 (left-handed 5-fold) have high docking scores at around the axes of the 5-fold (72^°^ in the figure) and 2-fold axis (180^°^). Among these two axes with high docking scores, the 5-fold axis was confirmed to be indeed stable based on RMSD, implying the consistency with the classification with the spherical tiling. This consistency was also confirmed for the other left-handed 5-fold capsids, 2BUK and 4V4M (Supplementary Figs. 6 and 7). 5ZJU (right-handed 3-fold) does not have stable docking structures at around the 5-fold axes. Instead, it has stable structures around the 3-fold axis (120^°^ in the figure) and the 2-fold axis. The 3-fold axis was confirmed to be indeed stable from the result of RMSD. Additionally, 1STM, classified as right-handed 2-fold, exhibits a single docking score peak at around the 2-fold axis. The structure around the 2-fold axis is consistently validated by the RMSD result.

7ODW, which is classified as left-handed and 5-fold, shows an exceptional result (Supplementary Figs. 6 and 7). In this case, we could not find notable docking score peaks except for the 2-fold axis. This could be interpreted as being due to 7ODW having interactions where subunits stack on top of each other in a direction perpendicular to the spherical surface (Supplementary Fig. 2), which would be not adequately characterized by 2D tiling. Also, 1M1C, which treats two subunits as one tile, shows exceptional behavior. While our model classifies it as right-handed 5-fold, the docking simulation results indicate that the 3-fold and 5-fold axes are stabilized to a similar degree (Supplementary Figs. 6 and 7). This suggests that our model may be underestimating the contribution of interactions due to the concave shape formed around the 3-fold axis, which is exceptionally created by the combination of two units.

As for references, we further performed the same type of docking simulation for non-capsid structurtes: the tabacco mosaic virus subunit (PDB ID: 6R7M. Fig. 4d) that only has a 16 1/3 symmetry axis and an NMR structure of chymotrypsin inhibitor structure in solution (PDB ID: 2M99. Supplementary Figs. 6 and 7) that does not have any symmetry axes. Our docking simulation correctly captured the stable symmetry axis for tabacco mosaic virus and do not show notable peaks except for 2-fold axis in the case of chymotrypsin inhibitor.

These results imply that during the initial stage of capsid self-assembly, there is a preference for subunits to form dimers along either the 3-, or 5-fold axes. In the theory of the nucleation-and-growth mechanism, subunits initially encounter to transiently form a dimer, which grows until it reaches a critical nucleus. Upon reaching this critical nucleus, the process enters a stable growth phase, progressively adding subunits until the complete capsid is formed. Whether the 3-fold axis or the 5-fold axis serves as the interface for the formation of this critical nucleus presents an intriguing question. Generally, the 5-fold axis is considered a candidate for the critical nucleus due to the greater stability resulting from increased inter-subunit contacts. However, the timescale of nucleus formation is 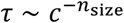 (where *n*_*size*_ is the size of the nucleus), making it kinetically unfavorable^1^. While the -3-fold axis has fewer interactions, it could provide a kinetic advantage or trade-off, suggesting the possibility of forming a trimer-based critical nucleus.

## Discussion

In this study, we have proposed a novel framework based on spherical tiling theory to explicitly describe the possible 2D shapes and interfaces of subunits in icosahedral capsids. This approach has allowed us to classify *T* = 1 real capsid structures in terms of subunit shapes and interfaces. Our findings reveal distinct interface groups even within the same *T* = 1 classification, highlighting variability in interaction patterns around 3-fold and 5-fold symmetry axes. Although we focused on *T* = 1 capsids due to their simplicity and the uniformity of interfaces across all subunits, the framework for describing subunit shapes proposed is not limited to *T* = 1 capsids. It can be naturally extended to capsids with other triangulation that follows CK theory, that is, when *T* = *h*^2^ + *hk* + *k*^2^ for integers (*h, k*) (see Supplementary Fig. 1 and text).

We believe that an explicit description approach of tile shapes and connectivity leads to a new theory of capsid structure that falls outside of CK theory. For example, in the *T* = 2 case, even though the shape of each tile is nearly identical, the connectivity between the tiles is not unique. Non-CK tiling is described as non-isogonal monogonal tiling on a sphere. In our study of 1M1C, we used the dimer as the subunit to be assembled. Since each subunit consists of two identical tiles, the whole system comprises a *T* = 2 assembly. This suggests that analysis and design of nested tiles, that is, the division of a single tile further into multiple identical units, may lead to a new way of understanding non-CK cases, such as *T* = 2,5,6, ….

The explicit description of subunit shape developed in this study also assists in the rational design of icosahedral protein complexes. In the design of icosahedral complexes, it is typical to first optimize the interactions either around the 3-fold or the 5-fold symmetry, and then stabilize the interactions around the remaining symmetry axes and the 2-fold axis. Generally, the choice of which symmetry axis (3-fold or 5-fold) to prioritize is not obvious. However, employing the method of our study allows for the determination of whether the 3- or 5-fold axis is more likely to stabilize from a given monomer structure. Furthermore, our approach enables the proposal of specific subunit shapes that satisfy spherical tiling for each symmetry axis. These tiling shapes can then be targeted for backbone design and sequence optimization using diffusion model-based frameworks, such as RFDiffusion^31^ and Chroma^32^, potentially enhancing the design process of icosahedral complexes.

Finally, we discuss the limitations of the proposed framework. The inherent constraint of this framework, based on spherical tiling, is limited to only 2D subunit shapes. There are cases in capsid structures where the shell thickness formed by the subunits is substantial relative to its radius, and the contributions from the three-dimensional interactions at the interface cannot be ignored, e.g., PDB ID of 7ODW in this study. In such cases, the stability of complexes likely affected by not only by the two-dimensional shape but also by three-dimensional shape complementarity between neighboring subunits. For these cases, an extension of the framework would be necessary, such as considering spherical tilings on spheres of various radii and addressing the cumulative effects of these tilings.

## Methods

### Data set of viral capsid structures

The capsid structures of *T* = 1 number were manually curated and selected according to the seqeuence variations and the experimental resolutions from VIPERdb version 3^34^. The selected structures include the satellite tobacco necrosis virus (PDB ID 2BUK^17^ and 4V4M^18^), the porcine circovirus 2 (3R0R^21^, 5ZJU^22^), the faba bean necrotic stunt virus (6S44^19^), the Haliangium ochraceum encapsulin (7ODW^20^), the satellite Panicum Mosaic Virus (1STM^23^), Sesbania mosaic virus deletion mutant (1VB4^24^). Also, the structure *T* = 1 number comprising 120-homomers was taken from the L-A virus (PDB ID 1M1C^25^). For use as a control reference, two non-capsid structures were also used. One is the tobacco mosaic virus (PDB ID 6R7M^35^ that has a lockwasher shaped ring with 16 1/3 subunits per turn. The other is the chymotrypsin inhibitor (PDB ID 2M99^36^) which is supposed to exist as a monomer in the physiological condition, lacking any symmetries in interacting modes with other monomers.

The subsequent analysis relies on the atomic coordinates of capsid structures obtained from the VIPERdb database. These coordinates have been pre-aligned to conform to a standardized icosahedral convention, known as the VIPER convention^37^, which ensures consistency in the orientation of the capsid structures. In this convention, two icosahedral 2-fold axes coincide with the *z* and *x* coordinate axes, while 3-fold and 5-fold axes lie between the *z* and *x* axes in the *xz* plane.

### Fitting of dihedron-unfolding model to experimental capsid structures

We here describe an optimization method to find a junction point that approximates the shape of the capsid from its point data, the radii of amino acids, and the symmetry axes. First, to compare the (normalized) 3D coordinates of the Cα atoms and tiles on a sphere, we use their gnomonic projections (from the center to *z* = 1 plane) ***p***_*i*_ and *R*_*di*_, respectively. First, we judged the chirality (left- or right-handed) by computing the maximum of minimum distances from the points (the projected positions of the Cα atoms) to the projected axes 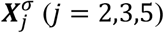, where *σ* = +1, −1 represents the left- or right-handed chirality.

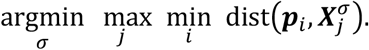

We create a region *R*_*ca*_ as the union of a circle of the average radius *r* of amino acid centering ***p***_)_. Then, we maximize the Dice coefficient between *R*_*ca*_ and the unfolded dihedron *R*_*di*_ (*x, y*) computed from the junction point *x, y*.

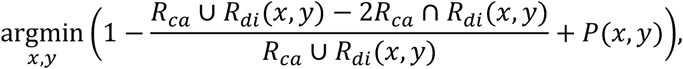

where *P*(*x, y*) is the penalty function to avoid self-intersection and to keep the spherical polygon inside a hemisphere. We used Genetic Algorithm solver *Galapagos*^38^ on the 3D CAD Rhinoceros/Grasshopper.

### Pairwise docking simulations and analysis of screw angles

Pairwise rigid-body docking simulations were performed for a pair of subunits taken from the whole capsid structure with ZDOCK^28,29^. Two subunits have identical structure with each other, thus structures obtained with the simulations are homo-dimers. At first, we applied ZDOCK with default settings, which outputs 2,000 top docking structures at a rotational sampling of 6^°^ interval, corresponding to 54,000 rotations. However, this setting did not yield sufficient statistics. Thus, we effectively performed dense rotational sampling by conducting 1,000 independent docking simulation runs with different initial seeds (used for the randomization of the orientations of initial structures). Finally, by sorting the results of independent docking runs, we obtained top 100,000 docking structures, which were then used for the subsequent screw motion and root mean square deviation (RMSD) analysis.

A spatial displacement of a rigid-body can be represented by a rotation about an axis and a translation along the same axis, which is called a screw motion. Here, to investigate the symmetry of the docking structures, we analyzed the structures in terms of screw motions. The screw axis and translation and rotation around that axis were determined by using the Rodriguez equations. Let ***p***_1_, ***q***_1_, and ***r***_1_ be position vectors of three atoms of monomer 1, and ***p***_2_, ***q***_2_, and ***r***_2_ be position vectors of three atoms of monomer 2. Then, the Rodriguez equations (http://robotics.caltech.edu/wiki/images/f/f3/Rodriguez.pdf) are written by,

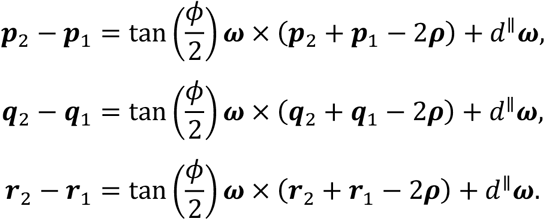

Here, ***ω*** is a unit vector parallel to the screw axis, ***ρ*** is a vector to a point on the screw axis, *ϕ* is the angle of rotation about the screw axis, and *d*^∥^ is the translation along the screw axis.

The RMSDs of the docking structure were evaluated by using the experimental structure as a reference. In the case of *T* = 1 structures, the number of picking a pair structure from a capid is □_60_*P*_2_ = 60 × 59 = 3,540 permutations. We calculated the RMSDs of individual docking structure to 3,540 reference structures, and plotted the minimum RMSD value at each region in the space of translation and rotation of the screw axis.

## Data availability

Codes and input files to reproduce the results of this paper are publicly available at https://github.com/matsunagalab/capsid. Large-size output files are available upon reasonable request to the corresponding authors.

## Acknowledgements

This work was supported by JSPS KAKENHI (Grant numbers: 20K21380, 22H04954 and 23K18105), and partly supported by MEXT as “Program for Promoting Researches on the Supercomputer Fugaku” (Development and application of large-scale simulation-based inferences for biomolecules JPMXP1020230119).

## Author contributions

T. Tachi conceived the idea of unfolding dihedrons for describing subunit shapes. R.T., S.N., T. Tendo, T.H., T. Tachi and Y.M. performed the research. R.T., S.N., T. Tachi and Y. M. wrote the manuscript. T. Tendo and T. H. edited the manuscript. All authors read and approved the final manuscript.

## Additional information

Supplementary Information accompanies this paper.

### Supplementary Information Note

**Supplementary Movie 1**. Unfolding of a dihedron when the junction point is inside the triangle.

**Supplementary Movie 2**. Unfolding of a dihedron when the junction point is outside the triangle.

**Supplementary Movie 3**. Unfolding of a dihedron. Cutting along a complex curve.

## Competing interests

The authors declare no competing interests.

## Notes

### Competing Interest Statement

The authors have declared no competing interest.

