## Supplementary Information for "Explicit description of viral capsid subunit shapes by unfolding dihedrons"

#### Description of subunit shapes of $T \neq 1$ capsid structures

The proposed method can be naturally applied to $T \neq 1$ in combination with the Caspar-Klug (CK) theory (Supplementary Fig. 1). In the CK theory, a regular triangle grid is converted to the grid on a regular icosahedron by converting a set of degree-6 grid points specified by two integers $(h, k)$ into degree-5 vertices. Instead of directly constructing spherical tiling, we first create planar tiling on the regular triangle grid[1]. Specifically, we first take out the triangle with angles 30°, 60°, and 90° consisting of one-sixth of the regular triangle, construct its double covering, and then cut out the dihedron and use its unfolding as a tile. This constructs a tiling under the $p6$ wallpaper group symmetry. These tiles can be applied to the deltahedron with $20 \times T$ ($T = h^2 + hk + k^2$) equilateral triangles to form a surface tessellation with $60 \times T$ tiles as desired.

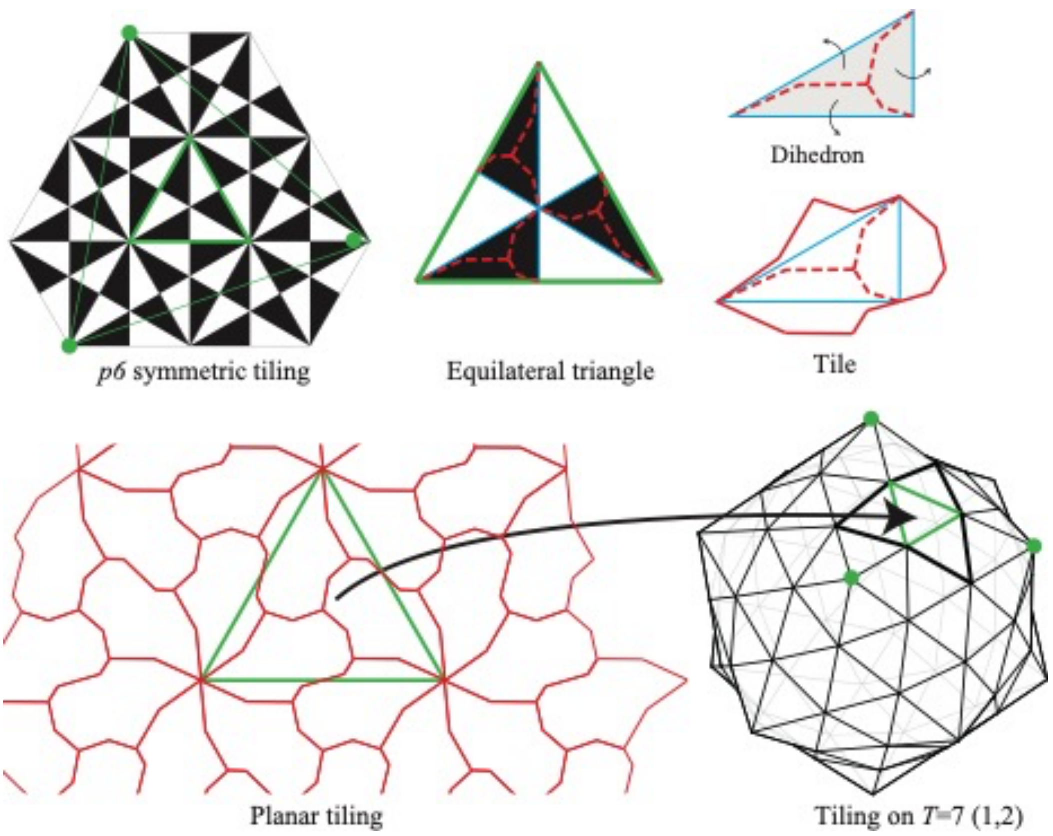


**Supplementary Figure 1.** Description of arbitrary subunit shapes of $T \neq 1$ capsid structures.

### Capsid structure data set

The capsid structures of $T = 1$ used in this study are summarized in Supplementary Fig. 2. These include the satellite tobacco necrosis virus (PDB ID 2BUK and 4V4M, the porcine circovirus 2 (3R0R, 5ZJU), the faba bean necrotic stunt virus (6S44), the Haliangium ochraceum encapsulin (7ODW), the satellite Panicum Mosaic Virus (1STM), Sesbania mosaic virus deletion mutant (1VB4). Also, the dimer structure of $T = 2$ number comprising 120 homomers was taken from the L-A virus (PDB ID 1M1C). For use as a control reference, two non-capsid structures were also used. One is the tobacco mosaic virus (PDB ID 6R7M that has a lockwasher shaped ring with 16 1/3 subunits per turn. The other is the chymotrypsin inhibitor (PDB ID 2M99), which is supposed to exist as a monomer in the physiological condition, lacking any symmetries in interacting with other monomers.

Sequence variations are analyzed using multiple sequence alignment (MSA). The MSA results are visualized by a phyrogenic tree in Supplementary Fig. 3. 2BUK and 4V4M share the identical sequence, and 3R0R and 5ZJU are almost identical with each other. The other sequences have low identities with each other.

| | Structure | PDB ID | Family | Genus | Resolution (Å) | Description |
|---|---|---|---|---|---|---|
| Left hand 5-fold | 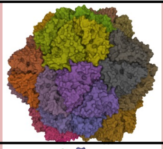 | 2BUK | Unclassified | Albetovirus | 2.45 | Satellite tobacco necrosis virus (STNV) |
| Left hand 5-fold | 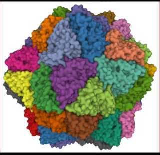 | 4V4M | Unclassified | Albetovirus | 1.45 | 1.45 Angstrom structure of STNV coat protein |
| Left hand 5-fold | 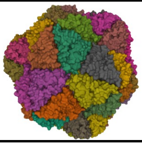 | 6S44 | Nanoviridae | Nanovirus | 3.19 | Faba bean necrotic stunt virus (FBNSV) |
| Left hand 5-fold | 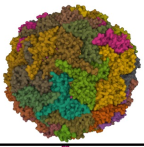 | 7ODW | Nanoparticles | Nanoparticles | 2.50 | Model of Haliangium ochraceum encapsulin from icosahedral single particle reconstruction |
| Right hand 3-fold | 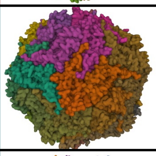 | 3R0R | Circoviridae | Circovirus | 2.35 | The 2.3 A structure of porcine circovirus 2 |
| Right hand 3-fold | 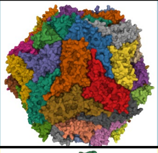 | 5ZJU | Circoviridae | Circovirus | 2.80 | Crystal structure of in vitro expressed and assembled PCV2 V particle |
| Right hand 2-fold | 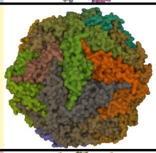 | 1STM | Unclassified | Papanivirus | 1.90 | Satellite Panicum Mosaic Virus |
| Right hand 5-fold | 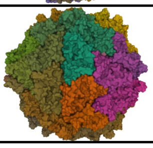 | 1VB4 | Solemoviridae | Sobemovirus | 3.30 | Sesbania mosaic virus deletion mutant Cp-N(δ)36 |
| Right hand 5-fold | 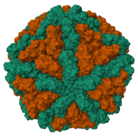 | 1M1C | Totiviridae | Totivirus | 3.50 | L-A virus |
| | 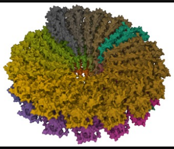 | 6R7M | Virgaviridae | Tobamovirus | 1.92 | Tobacco mosaic virus (TMV) |
| | 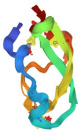 | 2M99 | NA | NA | NMR | Solution structure of a chymotrypsin inhibitor from the Taiwan cobra |

**Supplementary Figure 2.** Summary of PDB structures of capsids used in the study. They are classified according to the fitting results of the junction points (indicated by the background colors).

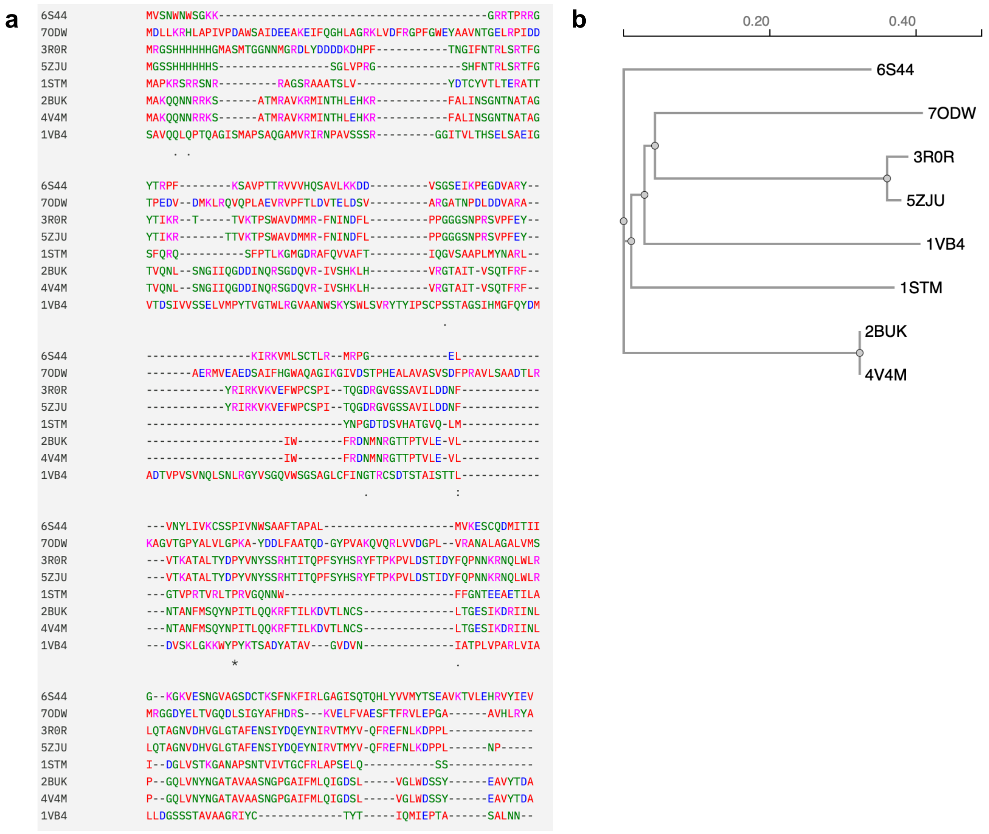


**Supplementary Figure 3.** Multiple sequence alignment and Phyrogenic tree of subunit sequences (except for 1M1C). Prepared with Muscle[2] on the EMBL-EBI job dispatcher sequence analysis tools framework[3].

### Junction points of subunit shapes

The coordinates of the junction points fitted to the experimental capsid structures and their corresponding subunit shapes are shown in Supplementary Fig. 4. As described in the text, the orientation (left- and right-handed) is determined by letting the boundary of the tile pass through the symmetry axes, while the coordinates of the junction points are determined by maximizing the overlap (Dice coefficient) between the subunit shape (orange in the figure) and the unfolded dihedron in the gnomonic-projection space.

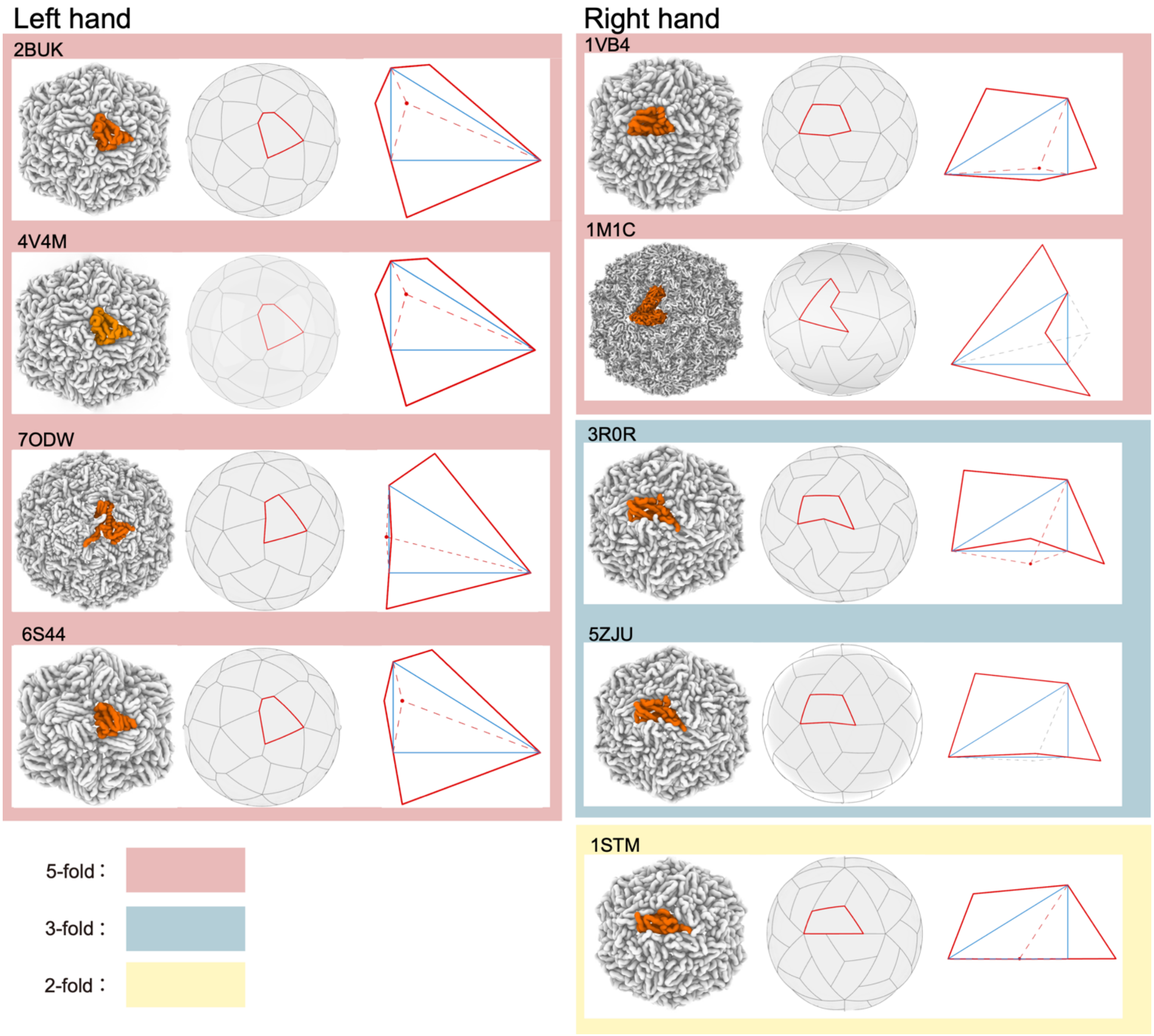


**Supplementary Figure 4.** Coordinates of the junction points and their corresponding subunit shapes. Classified according to the fitting results of the junction points (indicated by the background colors).

### Screw axes of subunit dimers

Supplementary Fig. 5 shows the screw axes calculated from the docked dimers of capsid subunits. As described in the main text, a screw axis describes a rigid-body motion in terms of translation and rotation along its axis.

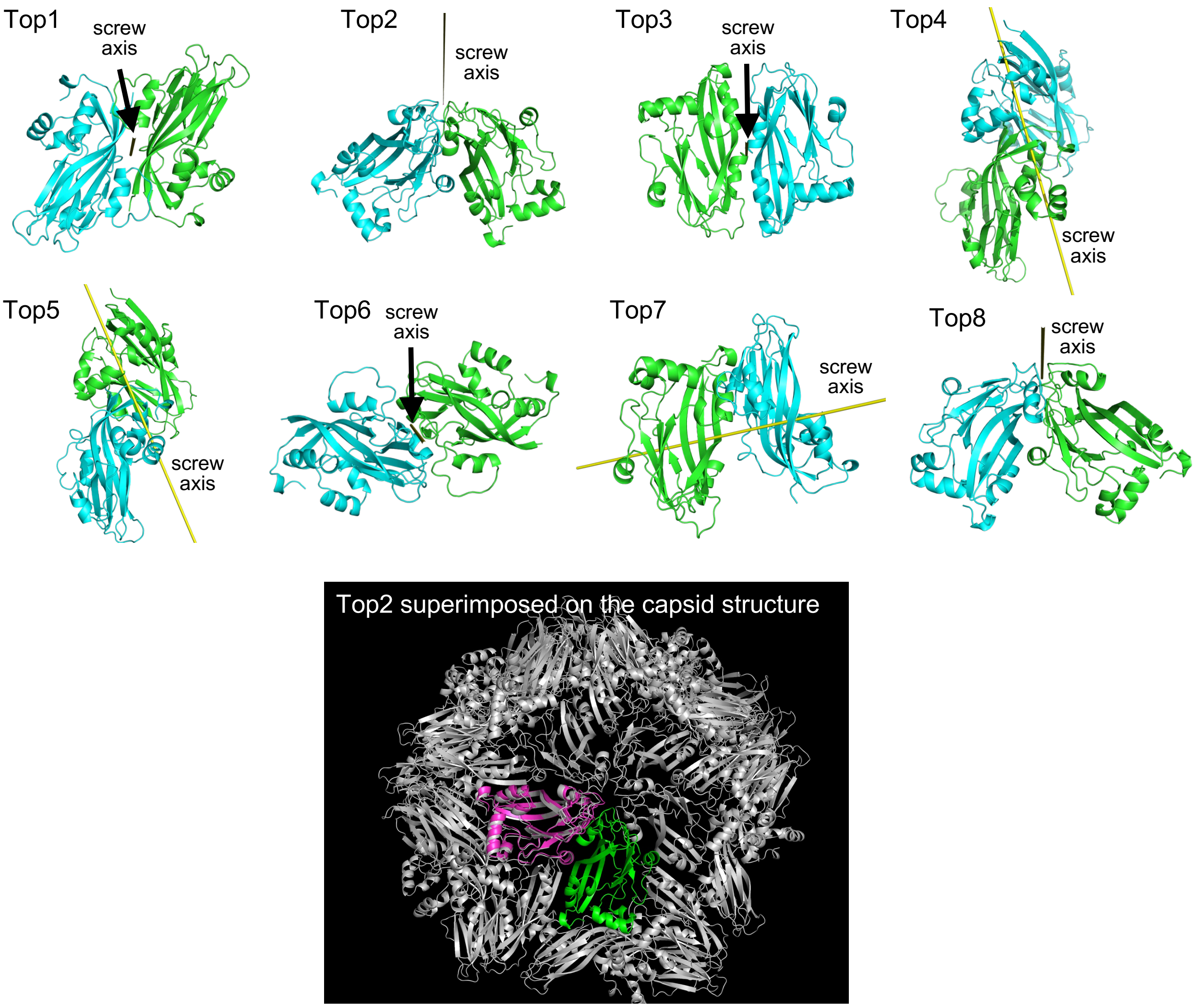


**Supplementary Figure 5.** Docking poses of paired subunits (PDB ID: 1VB4) with highest docking scores and their screw axes.

### Docking scores and RMSDs

Supplementary Fig. 6 shows the heatmap of the maximum docking scores of docked poses in the space of rotation and translation of detected screw motions. The figure show that the 2BUK, 4V4M, and 6S44 have high docking scores at around the rotation of 72°, and 120°, indicating that the stabilities of the interfaces of these symmetry axes. Contrary, 3R0R and 5ZJU have high docking scores at around 120° (3-fold axes). Supplementary Fig. 7 shows the heatmap of the minimum RMSDs of docked poses.

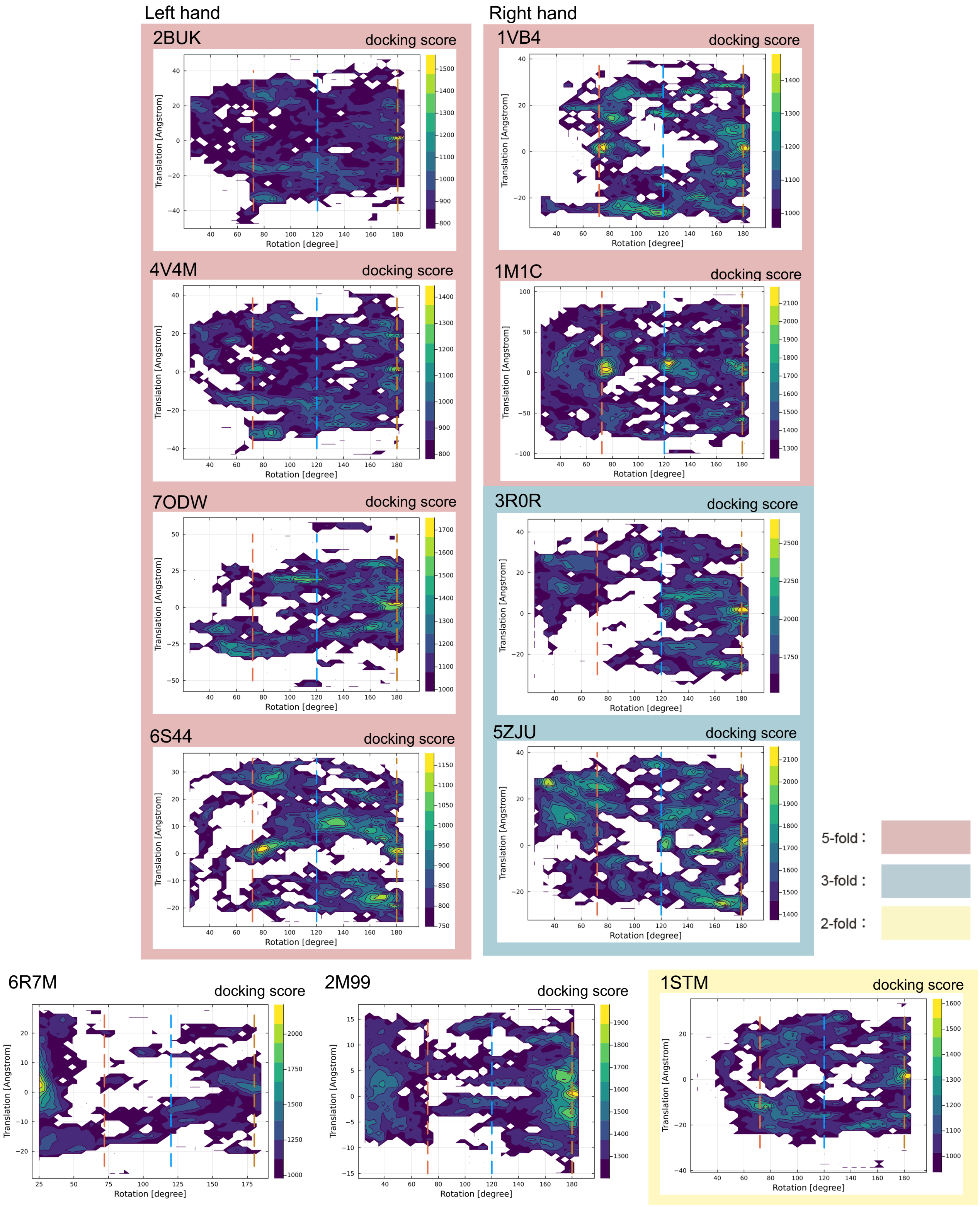


**Supplementary Figure 6.** Heatmap of the maximum docking scores of docked poses in the space of rotation and translation of detected screw motions. Dashed lines indicate the rotation positions of 72° (red), 120° (blue), and 180° degrees (ocher), which correspond to the 5-fold, 3-fold, and 2-fold axes, respectively.

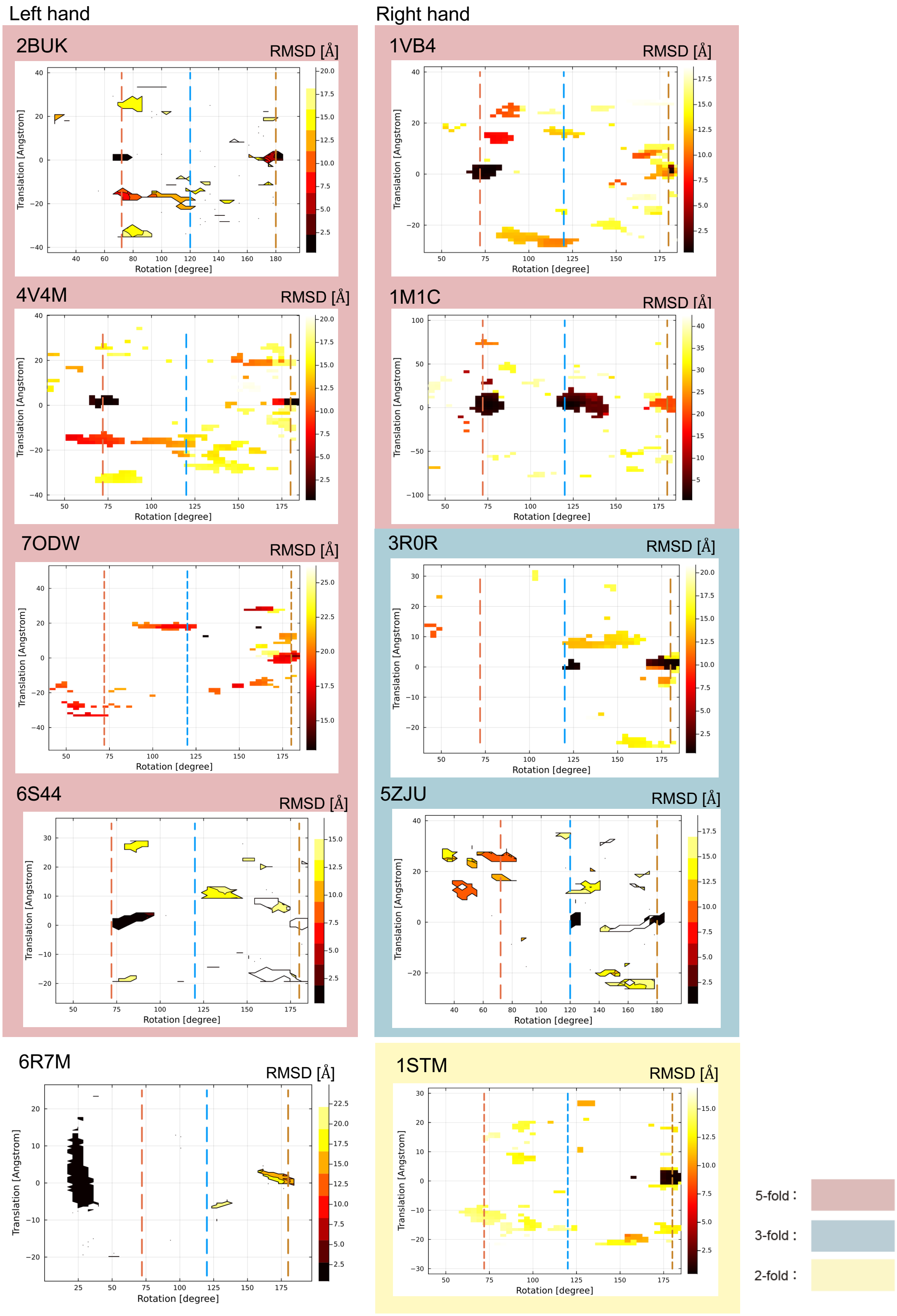


**Supplementary Figure 7.** Heatmap of the minimum RMSDs of docked poses from the experimental structure in the space of rotation and translation of detected screw motions. Dashed lines indicate the rotation positions of 72° (red), 120° (blue), and 180° (ocher), which correspond to the 5-fold, 3-fold, and 2-fold axes, respectively.
